# Comparative In-Vitro Physicochemical Evaluation and Dissolution Profiling of Glibenclamide 5mg tablet Marketed in Addis Ababa, Ethiopia

**DOI:** 10.1101/860163

**Authors:** Selass Kebede, Habtamu Abuye, Woldemichael Abraham, Sultan Suleman, Sileshi Belew

**Author notes:** Selass Kebede: (SK).

## Abstract

The safety of medicines is an essential part of patient safety. Global drug safety depends on strong national systems that monitor the development and quality of medicines. Poor quality medicines do not meet official standards for strength, quality, purity, packaging and labelling. Hence, this study determines *in-vitro* quality attributes of glibenclamide 5mg tablet marketed in Addis Ababa according to USP-38 drug monograph specifications. All tested brands meet the requirements for physical inspection & complied specification for friability and hardness. Besides, the tested brands met USP 38 specification for assay (99.96% to 108.85%) and for content uniformity (AV values ranges from 3.35 to 10.04). *In-vitro* release tests were carried out in phosphate buffer of 7.5 and 8.5 pH and showed drug release of ≥ 75%, met USP 38 requirements. However, significant difference with respect to dissolution profile among tested brands GL4 and GL6 were confirmed with comparator product through model independent approach. Moreover, DE values were studied and confirmed that GL4 and GL6 were not therapeutically interchangeable with innovator product.

## Introduction

Glibenclamide, which is also Glyburide in USA, is a second-generation sulfonylurea oral hypoglycemic agent used in the treatment of noninsulin-dependent diabetes. It is a Biopharmaceutical classification system (BCS) class II drug that has high permeability and poor water solubility (Figure 1).[1] It is one of the most prescribed long-acting anti-hyperglycemic agents that lower the blood glucose acutely by stimulating the release of insulin from the pancreas, an effect dependent upon functioning beta cells in the pancreatic islets. [2]

**Figure 1.**
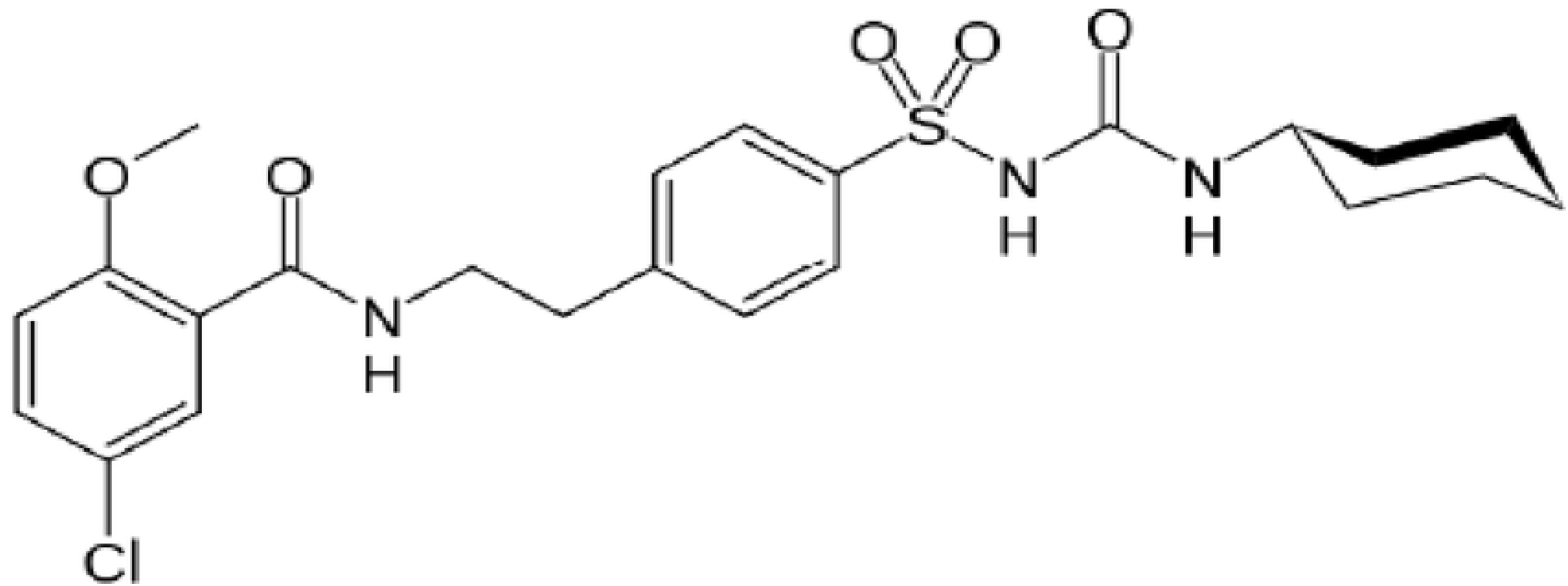
Chemical structure of Glibenclamide

The food and Drug Administration (FDA) has dictated that in order for the generic drugs to be approved, they have to pass many guided tests and examination for their physicochemical characteristics, contain the same active constituents and strength, in addition, to be bioequivalent to their innovator product. [3]

The quality concern of drugs is as old as drugs themselves. Despite all the advances made over the years, this concern has not disappeared. In the recent past, the unregulated proliferation of pharmaceutical industries and products has brought with it many diverse problems of varying magnitude. [4] The use of ineffective, poor quality, harmful medicines can result in therapeutic failure, exacerbation of disease, resistance to medicines and sometimes death. It also undermines confidence in health systems, health professionals, pharmaceutical manufacturers and distributors. Money spent on ineffective, poor quality medicines is wasted – whether by consumers or governments. [5]

The safety, efficacy, and quality of the medicines should be ascertained to provide a desired pharmacological effect. Substandard drugs have been defined as those which do not meet quality specifications set for them, as to contain under or over concentration of ingredients, contamination, poor quality ingredients, poor stability and inadequate packaging. [6]

Many studies have shown the presences of substandard drugs available in the markets of several countries and significant difference in the dissolution and disintegration of tablets were revealed. A study conducted in Jordan indicated that generic Glibenclamide products exhibits a significant difference in their dissolution profile when compared to innovator products and 60% (3 of 5) tested generics were found to be out of British pharmacopoeia specification for assay values. [7] Besides, a study done in Saud Arabia pointed out that 100% (5 of 5), 60% (3 of 5) and 20% (1 of 5) of tested Glibenclamide products fails to release ≥75% of their content at PH value of 7.4, 7.8 and 9.5 respectively after one hour. [8] Moreover, 13 of 142 Glibenclamide tablet formulation from 28 countries (from Europe, north and South America, Africa, Asia and Australia) showed marked differences in *in-vitro* dissolution behavior. [9]

Ethiopian Food, Medicine and Health care Administration and Control Authority drug registration guideline requires every generic product to have a valid bioequivalence study that demonstrates its equivalence to the originator brand. Few studies done on the country, however, revealed that circulation of low quality medicines in its jurisdiction. A nationwide study on anthelminthic and antiprotozoal preparations available in Ethiopia market reported that 45.3% of tested samples were unable to comply pharmacopoeias specifications. [10] Gebrezgabiher *et al* localized study also showed 16.67% menbendazol tablet failed for disintegration & dissolution test. [11] In the same vicinity, study conducted on ciprofloxacin indicated that 16.67% failed to release 80% within 30. [12] Additionally, Lantider k. *et al* reported 62.5% amoxicillin marketed in that area was not pharmaceutically interchangeable. [13]

This findings are few evidences that Ethiopian health system endangered with poor quality medicines circulation within its drug supply chain system. Hence, the objective of the present study was to determine the physicochemical quality and dissolution profiling of imported and locally manufactured Glibenclamide 5mg tablets marketed in Addis Ababa, Ethiopia.

## Materials and methods

### Materials

#### Chemicals

Reference standard of glibenclamide was obtained from Cadila Pharmaceutical Manufacturing private limited company (Addis Ababa, Ethiopia). High-performance liquid chromatography (HPLC) grade acetonitrile and methanol, potassium di-hydrogen orthophosphate, monobasic ammonium phosphate, sodium hydroxide, phosphoric acid & Di-ionized water were obtained from Jimma University Laboratory of Drug Quality-JuLaDQ (Jimma, Ethiopia).

#### Apparatus

The Agilent series HPLC system (Germany) employed in this study consisted of a Diode array/Ultra violet (UV) detector (Merck-Agilent, model Agilent 1260), a quaternary pump (Merck-Agilent, model Agilent 1260), an integrator unit (Merck-Agilent, model Agilent 1260) and equipped with chem. Station 32 software. The employed HPLC column was C18, 5μm particle size, 150mm length and 4.6 mm internal diameter (Thermo Scientific, USA). Dissolution experiments were carried out using a RC-6D Tian Jin scientific dissolution apparatus manufactured in China and Cecil Aquarius UV-Visible spectrometer (England). Friability testing was carried out using Pharma PTFE Friabilator (Germany). Hardness test was carried out using three in one Pharma PTB hardness tester. Measurements of pH were made using AD1020 PH/mv/ISE and Temperature meter (Hungary). Weighing was done by using METTELER TOLEDO analytical weight balance (Switzerland). Water purification was done by using water distiller and de-ionizer instrument.

### Methods

The In-vitro critical quality attributes of the glibenclamide 5mg tablets were assessed according to USP-38 monograph specification. [14] Besides, additional *in-vitro* quality control parameters were used to investigation the products.

#### Identification

The retention time of the major peak of the sample solution was compared with that of the standard solution as stated in the USP/NF 38 drug monograph. [14]

#### Assay and content uniformity

##### Standard preparation

10mg of working standard was weighed using verified analytical balance and put in to 25 ml volumetric flask. Acetonitrile equivalent to 20 ml was added and shacked. Then, water equivalent to 0.4 ml per mg of Glibenclamide was added and shacked for 30 minutes. [14] Then, the solution was used to check system suitability of the methods.

##### Sample preparation

Ten tablets from each sample were taken and placed individually in a suitable container. Water equivalent to 0.4 ml per mg of glyburide was added and swirled to disperse and wet tablet material. Then acetonitrile equivalent to 2.0 ml per mg of glyburide was added and shaken for 30 min. [14] The suspension was centrifuged and the clear supernatant was used.

##### Chromatographic system

The liquid chromatographic system equipped with a 254-nm detector and 4.6-mm × 150-cm column that contains packing L7 having flow rate of 2 ml per minute with 10 µl injection volume was used to carry out the assay measurement. [14]

The analysis result was computed with the following formula:

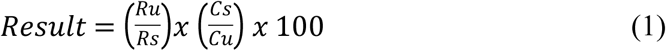

Where; R_u_ is response of unknown sample, R_s_ is response of reference standard C_s_ is concentration of the reference standard; C_u_ is concentration of unknown sample

#### Dissolution

The tests were performed according to pharmacopoeia specifications using Apparatus 2 (paddle method). The medium employed was 900 ml of 0.05M phosphate buffer of pH 7.5 and 8.5. Medium temperature was set at 37°C ± 0.2°C. Six tablets of each product were placed in the dissolution apparatus (one in each vessel). Samples (5 ml) were withdrawn at pre-determined time points (5, 15, 30, 45 and 60 at 7.5 pH; 5, 15, 30 and 45 at 8.5 pH) and the withdrawn samples were replaced with respective buffer solution. All samples were then filtered before being measured. HPLC system equipped with a 254-nm detector and 4.6-mm × 25-cm column that contains packing L7 for buffer medium of 7.5 pH and HPLC system equipped with a 215-nm detector and 4.6-mm × 25-cm column that contains packing L7 for buffer medium of 8.5 pH was used to carry out the dissolution measurement. The flow rate was adjusted at 2 ml per minute and with 75 µl injection volumes for 7.5 pH buffer medium and 1.5 ml per minute and with 50 µl injection volumes for 8.5 pH buffer medium. The systems were functioned with relative standard deviation of NMT 2.0% for replicate injection. [14] The amount of dissolved glibenclamide was determined using the regression equation computed with reference sample with the same buffer type.

The analysis result was computed using the following formula:

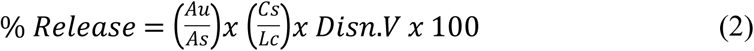

Where; A_u_ is peak response of the sample, A_s_ is peak response of the reference C_s_ is concentration of the reference, L_c_ is label claim, V is dissolution volume And, using regression equation:

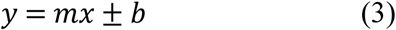

Where; m is the slope, b is the intercept

The dissolution pattern of the drug was analyzed using both model-dependent methods—zero order, first order, Weibull approach, Hugichi, Hickson-crowell and Korsmeyer-Peppas; and model-independent methods—fit factors were used to compare similarity and dissimilarity between tested brands and comparator. [15 & 16] Model-dependent method equations:

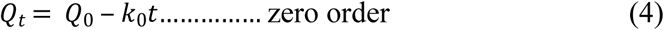

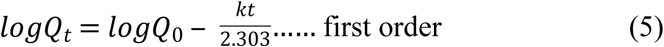

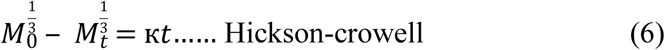

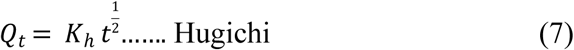

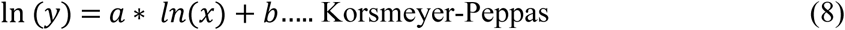

Model-independent method equations:

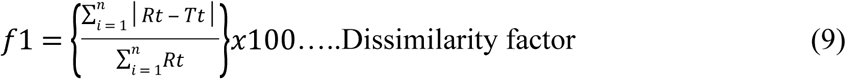

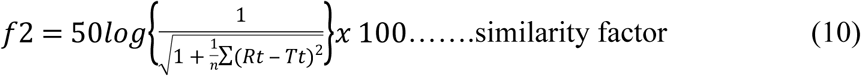

#### Friability

Twenty tablets from each brand were weighed individually and subjected to abrasion using a tablet friability tester at 25 revolutions per minute for 4 minutes. The tablets were weighed again and the difference in weight will be calculated as the percentage. A maximum weight loss of not more than 1% of the weight of the tablets being tested are not allowed.

The result of the test was computed using the following formula:

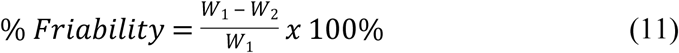

Where W_1_ is initial weight of randomly chosen 20 tablets, W_2_ is after subjecting the tablets to the friabilator for 4 minutes at 25 revolutions per minute, the final weight.

#### Uniformity of weight

Twenty tablets from each of the brand was weighed individually using analytical balance. The average weights of the tablets for individual brand and their deviation from the average weight was calculated.

And % of weight variation was calculated using the following formula:

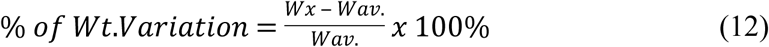

Where W_x_ is individual weight, W*_av._* is average weight tested tablets,

#### Crushing strength/Hardness

Hardness of individual brand drugs was obtained by measuring the crushing strength of ten randomly selected tablets using hardness tester. Then the mean and standard deviation was calculated.

### System verification

#### Method accuracy/recovery

As indicated in International Conference on Harmonization (ICH) guideline Q2B, system recovery was done by adding known amount of standard in the blank. The working standard has been prepared in three different concentrations (one from the lower, one from the middle and one from the higher amount) with in the range of 70% to 130% concentration. Each of the three concentrations (78%, 104% and 119%) has been injected in triplicated manner in to the HPLC system and % recovery for individual concentration has been computed. Since the difference in % recovery values at each concentration becomes less than 2% which is the maximum limit set in the ICH Q2B guideline, the system was proved to be fit for the purpose. [17] The result of recovery tests were presented in Table 1.

**Table 1.** System Accuracy/recovery test result at 3 different concentration (n=3).

| Theoretical value | % value | Measured value | % recovery | Difference in % recovery | Acceptance limit $\pm 2\%$ |
| --- | --- | --- | --- | --- | --- |
| 78 | 100 | 77.99 | 99.99 | 0.01 | pass |
| 104 | 100 | 103.93 | 99.93 | 0.07 | pass |
| 119 | 100 | 118.55 | 99.62 | 0.38 | pass |

#### Method Precision

The repeatability of the method was checked by using prepared working standards at three different concentrations of 90%, 100%, and 110% within the range of 80% to 120% as clearly stated in the ICH Q2B guideline. So, triplicate injections from each concentration were injected to the system and the RSD for individual concentration was computed. The calculated Relative Standard Deviation (RSD) values for 90%, 100% and 110% were 0.07, 0.05 and 0.20 respectively. Thus, the method is proved to be fit for the purpose since the RSD values are below acceptable maximum limit of 2 as indicated in ICH Q2B. [17] The result of the repeatability tests were presented in Table 2.

**Table 2.** System Repeatability test result at 3 different concentration (n=3).

| % Conc. | Peak | Peak Avg. | % Conc. | Avg. Conc. | SD | RSD | Acceptance limit $\pm 2$ |
| --- | --- | --- | --- | --- | --- | --- | --- |
| 90 | 1561.92 | 1560.75 | 90.39 | 90.33 | 1.02 | 0.07 | pass |
| 90 | 1560.04 |  | 90.29 |  |  |  |  |
| 90 | 1560.3 |  | 90.30 |  |  |  |  |
| 100 | 1818.98 | 1818.37 | 103.96 | 103.93 | 0.81 | 0.05 | pass |
| 100 | 1818.67 |  | 103.95 |  |  |  |  |
| 100 | 1817.45 |  | 103.88 |  |  |  |  |
| 110 | 1922.54 | 1918.27 | 109.43 | 109.21 | 3.75 | 0.20 | pass |
| 110 | 1916.74 |  | 109.12 |  |  |  |  |
| 110 | 1915.51 |  | 109.06 |  |  |  |  |

#### Method linearity

The linearity of the method has been verified by using five different concentrations within the range of 80% to 120%. Triplicate injection from each concentration has been injected to the system and RSD for individual concentration were computed. The system was found to be fit for the purpose as the calculated RSDs for all concentration become less than 2% which is the maximum acceptable limit stated in ICH Q2B guideline. [17] The result of linearity tests for assay and content uniformity method was indicated in Fig 2.

**Figure 2.**
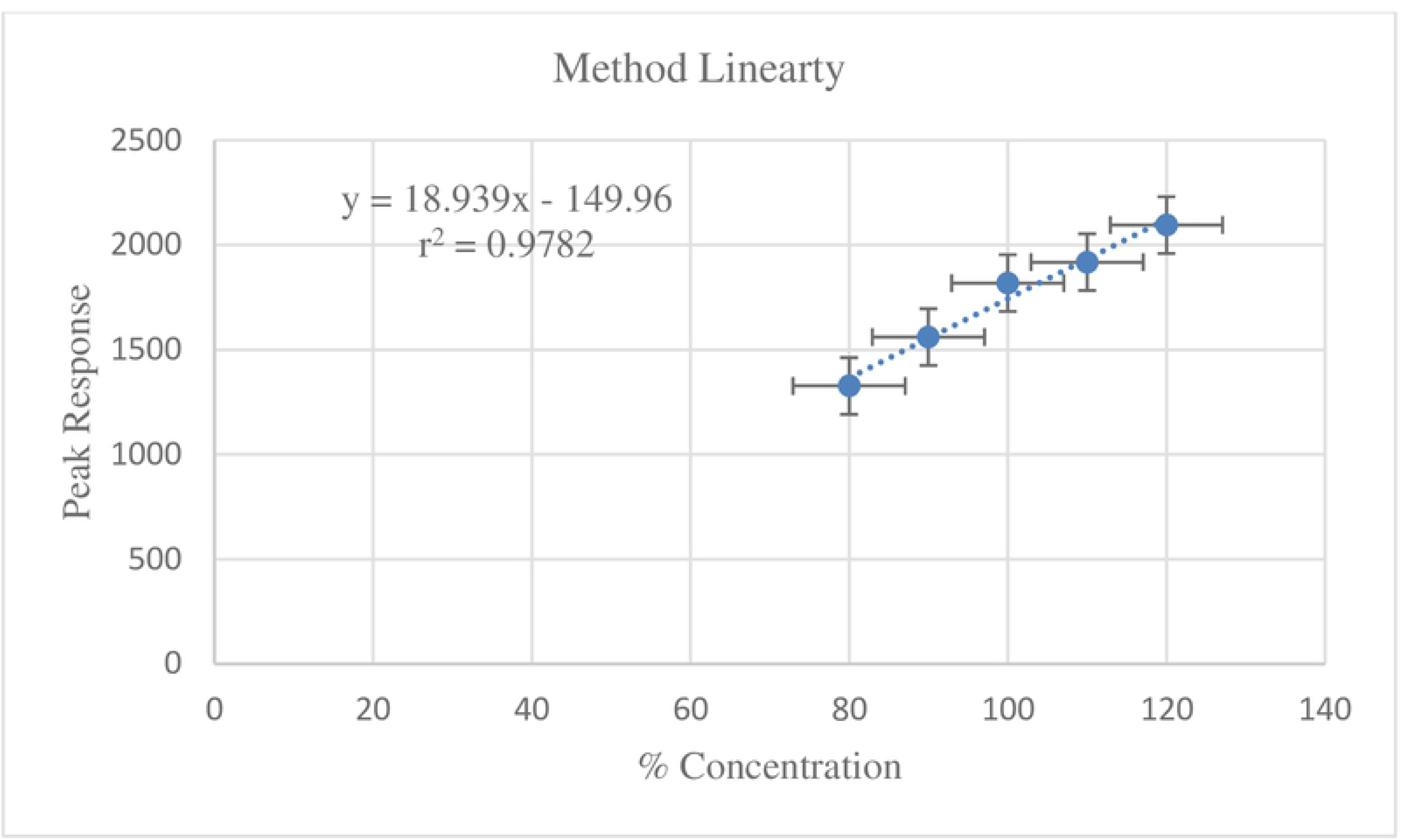
System linearity test result at 5 different concentration of the standard (n=3).

For a dissolution test, methods linearity was computed by using regression equation: y= mx ± b; where; m is the slope, b is the intercept. For buffer medium of 7.5 pH the regression equation was y= 0.008x + 0.0919 with regression coefficient (*r*^*2*^) of 0.9153 whereas at 8.5 pH buffer the regression equation was y= 0.004x + 0.2321 with regression coefficient (*r*^*2*^) of 0.9876. Thus, the method was found to be suitable for the analysis.

#### Statistical Analysis

Assessment of quality of each sample was performed according to the test method described in standard pharmacopeia (USP-38). The data generated were subjected to appropriate software’s such as excel-2010, Minitab and kinetic DS. In addition both model dependent and model independent approaches were used to compare dissolution profile between the Innovator and generic brands of glibenclamide 5mg tablet.

## Results and discussion

Glibenclamide 5mg tablet products were purchased from the local drug retail outlets/health facilities in Addis Ababa, Ethiopia in the same way that the patient might have bought them from such facilities by using mystery shoppers during the study period. The product information of glibenclamide 5mg tablet included in the study was presented in Table 3 with their full description.

**Table 3.**
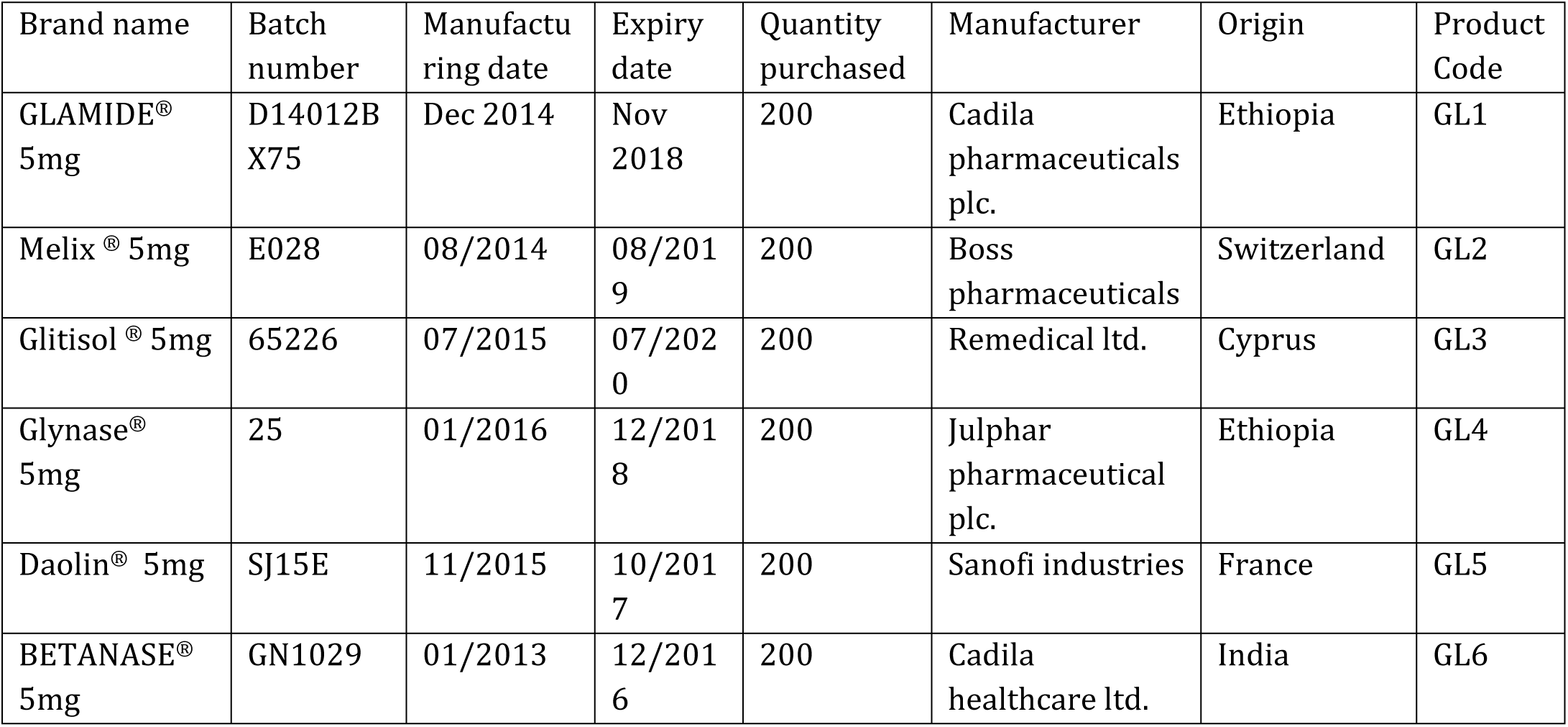
Product information of different brands of Glibenclamide tablet included in the study.

The physical characteristics of the product are a predetermining factor that influences the dissolution behavior and API content of the drugs. As a result, significant difference in the physical characteristics among different brands of a given drug results in great variation in the efficacy of the drug which leads to either treatment failure and/or toxicity to the patients. [9] As indicated in Table 4, tested glibenclamide brands comply for physical characteristics, packaging and labelling information requirements. Similar results have been reported for compliance in packaging and labeling in different countries. [7-9]

**Table 4.**
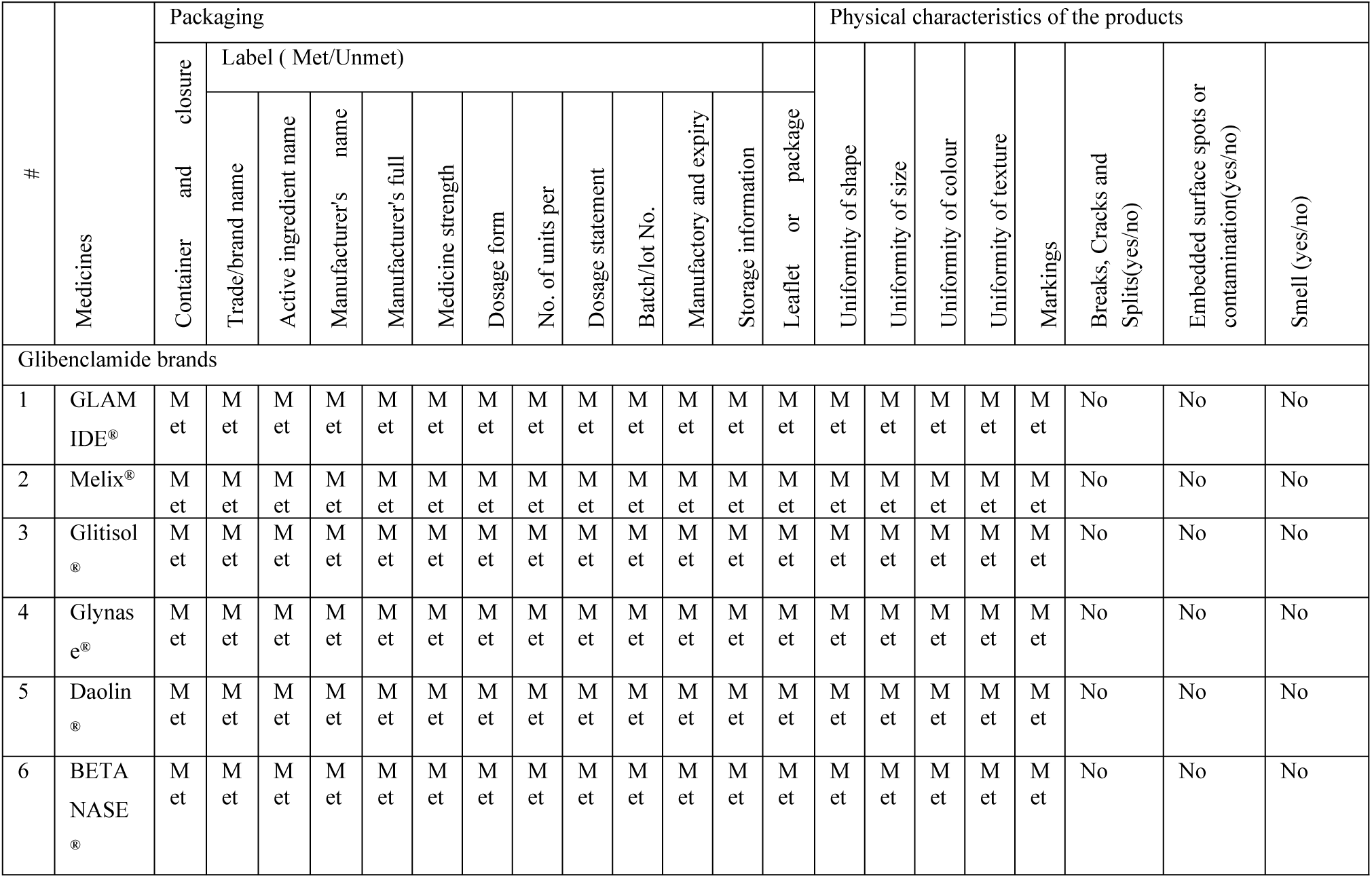
Physical characteristics and packaging and labelling information of samples of different glibenclamide medicines.

### Identification test

The identification test was conducted according USP 38 drug monograph specification. A reasonable peak shape was obtained for the analyte as per system suitability requirements. The identity of tested products was confirmed by comparing their retention time with that of glibenclamide working standard. [14] As indicated in Figs 3-5, tested glibenclamide products exhibited practically similar retention time (Rt) values when compared to reference standard. Thus, it can be concluded that all the tested glibenclamide products contains correct active ingredient.

**Figure 3.**
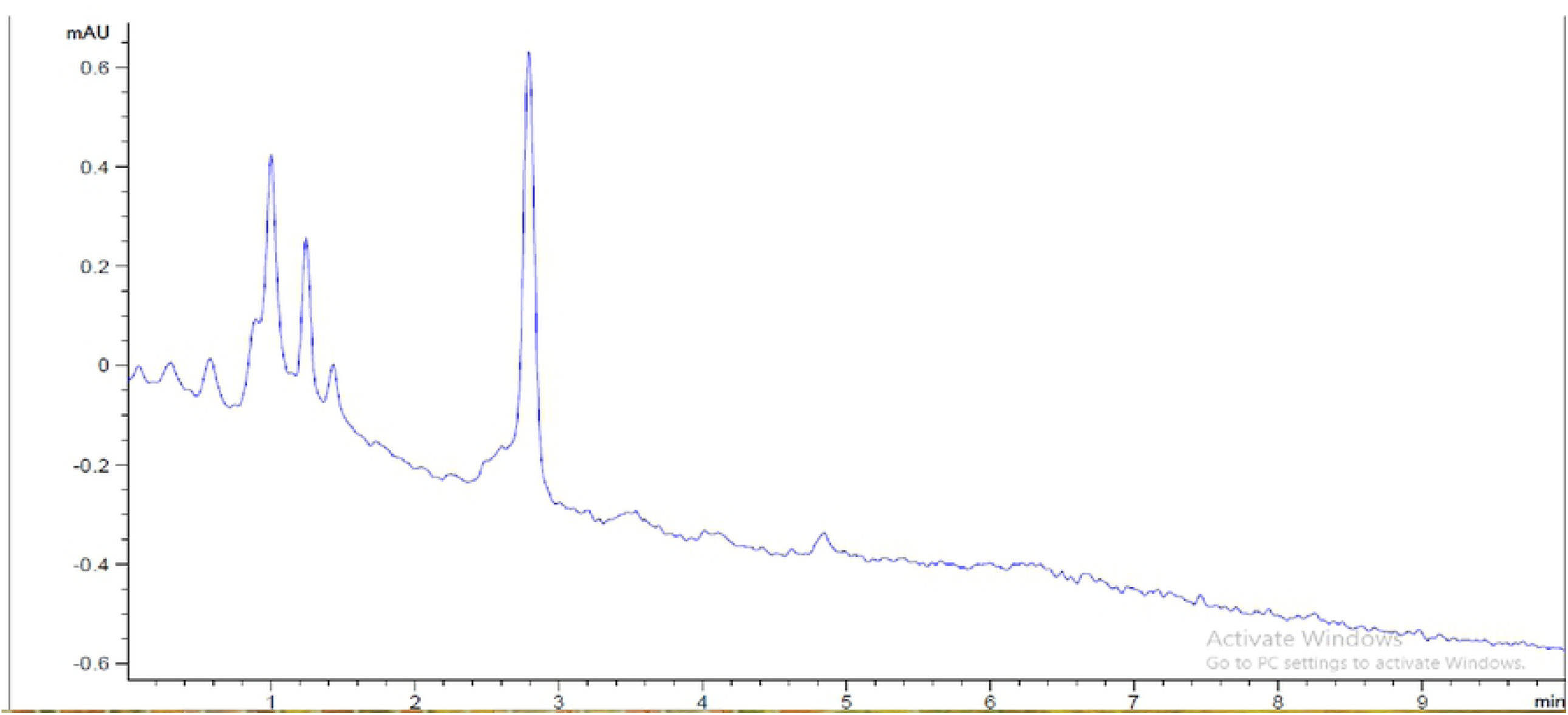
Blank chromatogram

**Figure 4.**
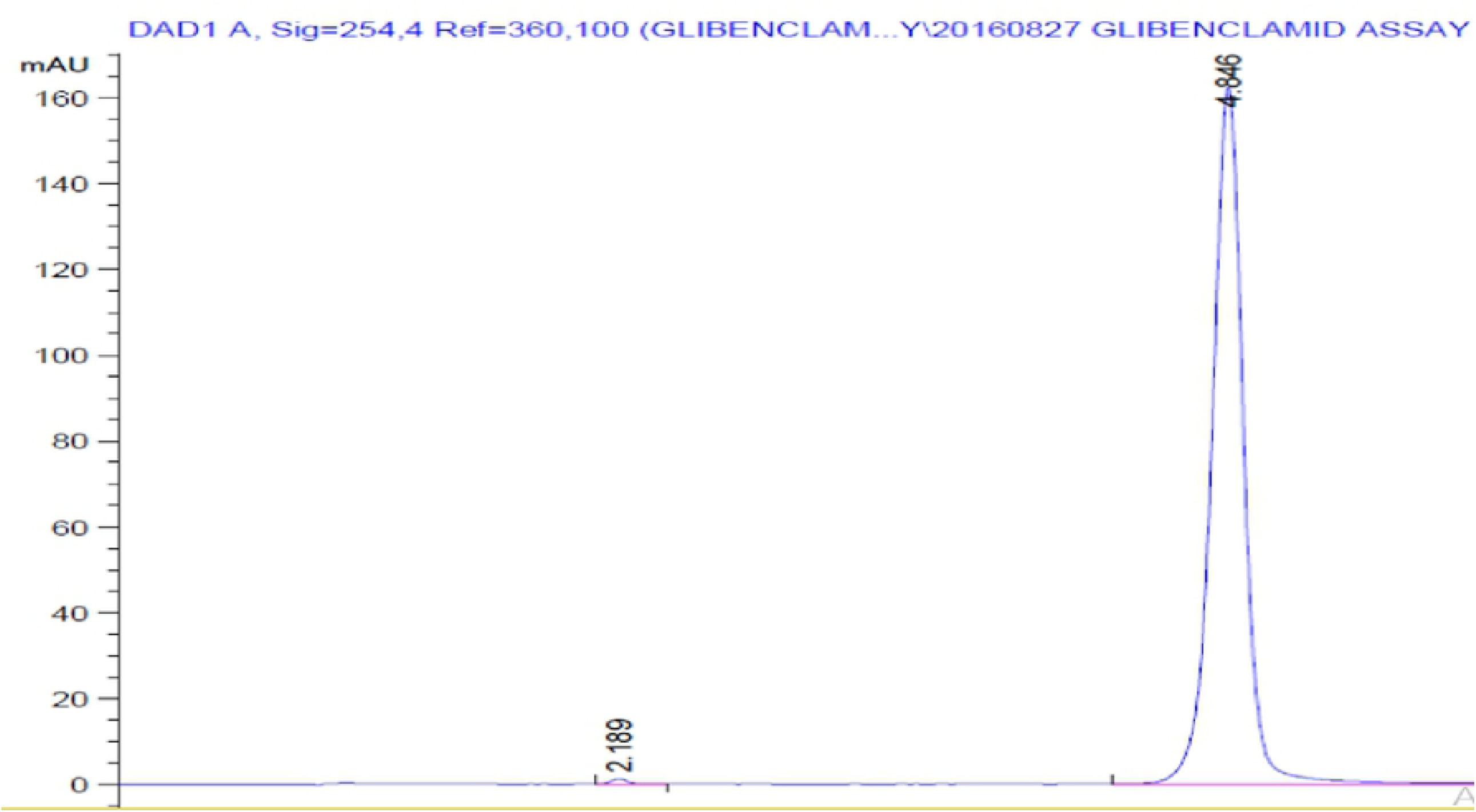
Reference chromatogram

**Figure 5.**
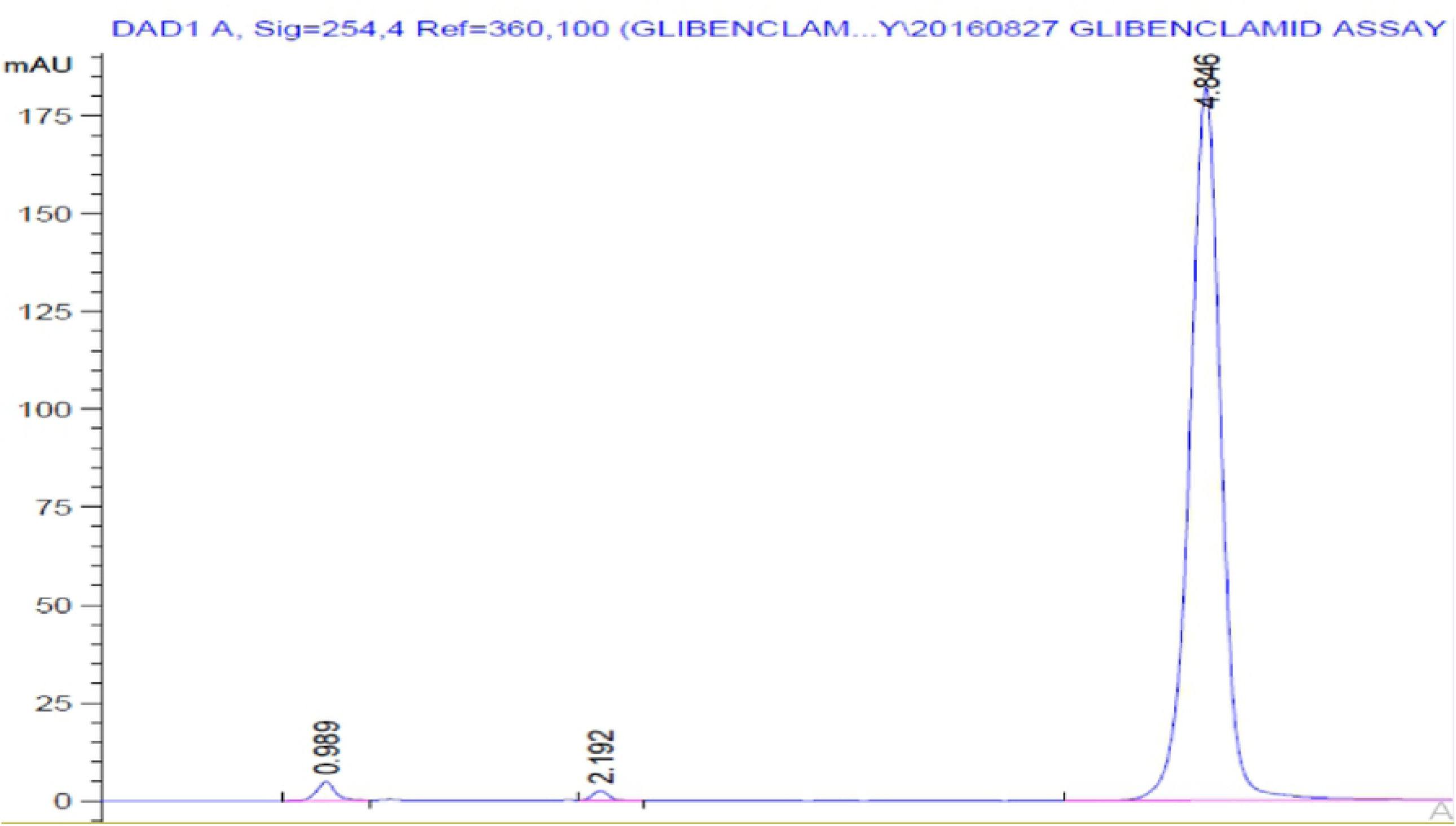
Sample chromatogram

### Assay for API

All six brands of Glibenclamide 5mg tablet were assayed as indicated in USP 38 monograph. [14] As presented in Table 5, the assay results of tested glibenclamide brands were found to be within the range of 99.97% to 108.85% which was within USP 38 assay specification (90% to 110%). However, brands GL1 and GL6 failed to pass British Pharmacopeia (BP) and European Pharmacopeia (Ph.Eur) specifications for assay (95% to 105%). There were reports of similar incidents that showed some of the generics were not in line with BP, Ph.Eur. and International Pharmacopeia (Ph. Int.) specification for their API contents. [7] Among all tested glibenclamide products, GL1 had higher assay value of 108.85% whereas GL5 had lower assay value of 99.97%.

**Table 5.** Average percentage per label obtained for each tested product.

| Product Name | Product code | Assay Avg. | RSD of assay | Reference Acceptance limit USP (90-110) |
| --- | --- | --- | --- | --- |
| GLAMIDE® 5mg | GL1 | 108.85 | 0.17 | Pass |
| Melix® 5mg | GL2 | 101.08 | 0.08 | Pass |
| Glitisol® 5mg | GL3 | 102.19 | 0.16 | Pass |
| Glynase® 5mg | GL4 | 101.05 | 0.12 | Pass |
| Daolin® 5mg | GL5 | 99.96 | 0.47 | Pass |
| BETANASE® 5mg | GL6 | 105.52 | 0.05 | Pass |

The difference in the assay values between tested generics and comparator product were assessed using annova single factorial analysis at 95% CI. The calculated P value and F value were found to be 6.41584E-21 (lower than expected value of 0.05) and 896.65 (higher than Fcrit value of 2.772853153) respectively. Both P and F values indicated as there is significant difference among assay value of tested brands. Post ANNOVA t-tests with Bonferroni correction were used to indicate the brands that have significant difference from the comparator. Thus, except GL4, the mean assay values of all generics were found to have significant difference with comparator product since the T stat value of GL1, GL2, GL3 and GL6 were 34.86, 4.64, 8.89 and 23.35 respectively which were greater than the T critical value of comparator GL5 (which was 2.31). This has been confirmed with calculated P value of the products that become less than the Bonferroni corrected expected P value at 95% CI.

### Content uniformity

The content uniformity test was performed as per USP 38 specification for all glibenclamide tablets and the acceptance value was calculated. [14] The acceptance values for all brands were found to be less than 15 (the maximum allowed acceptance value, L1) as shown in Table 6. Thus, it can be concluded that all brands comply USP-38 requirements for content uniformity.

**Table 6.** Content uniformity test result (n=10).

| Product code | Calculated acceptance value | Maximum allowed AV value L1=15 | Reference |
| --- | --- | --- | --- |
| GL1 | 8.42 | Pass | USP-38 |
| GL2 | 10.04 | Pass |  |
| GL3 | 5.78 | Pass |  |
| GL4 | 4.71 | Pass |  |
| GL5 | 3.35 | Pass |  |
| GL6 | 6.18 | Pass |  |

### Dissolution

Dissolution test is a critical quality attribute parameter used to predict the *in-vivo* performances of oral pharmaceutical solid dosage forms like tablets and capsules. Besides, it can serve as a surrogate for bioavailability and bioequivalence. [18 & 19] In this study, dissolution tests were conducted with paddle dissolution apparatus as specified in USP-38 monograph for glibenclamide tablet preparation using phosphate buffer as a dissolution medium at 7.5 and 8.5 pH values.

The dissolution profiles of tested products at 7.5 pH phosphate buffer, as indicated in Fig 6, were in line with pharmacopoeia requirements for percent drug release since all tested brands exhibits drug release behavior of more than 75%. Thus, it can be concluded that all brands have passed dissolution criteria.

**Figure 6.**
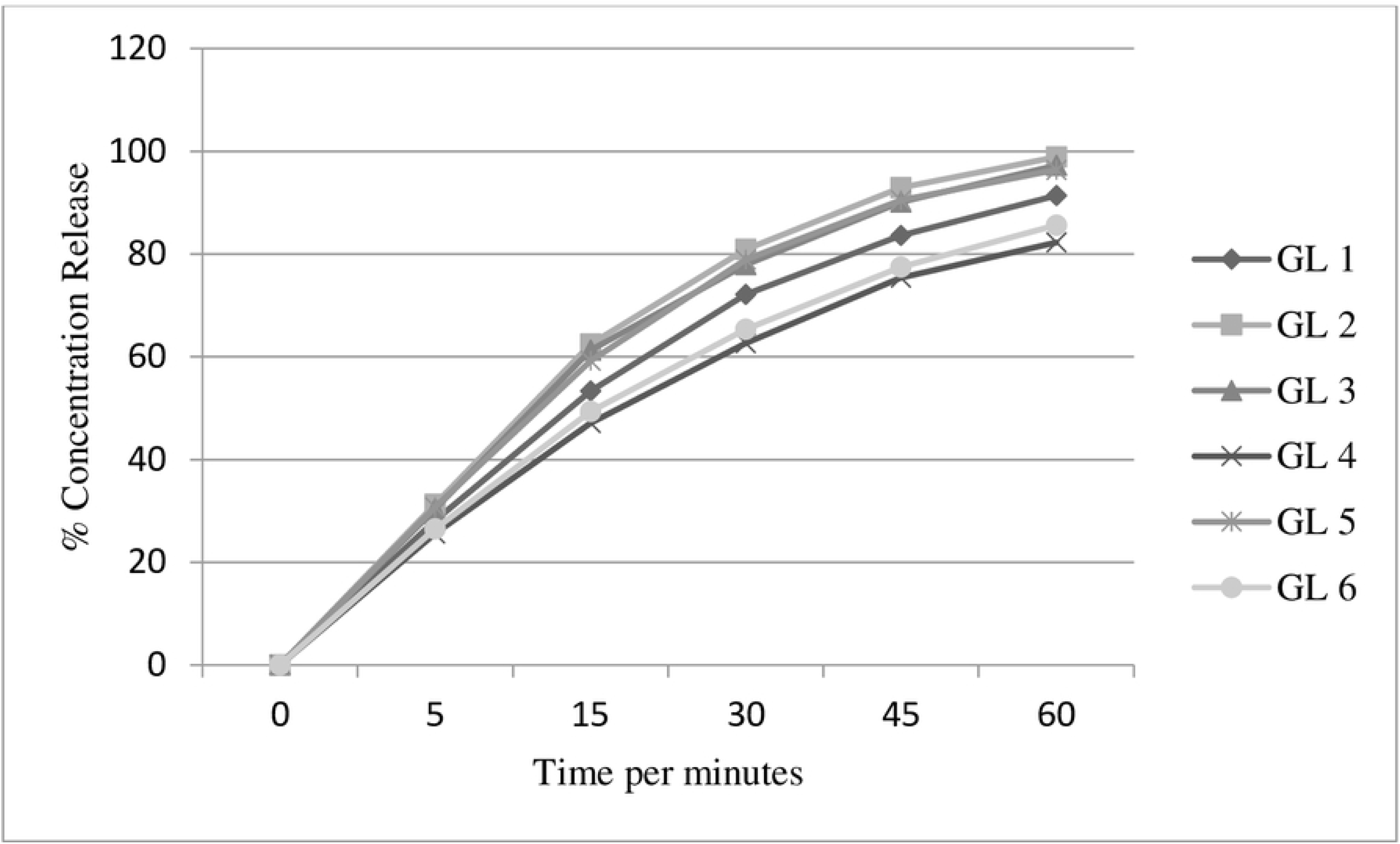
Dissolution profile of tested drugs at 7.5 pH phosphate buffer.

The two way ANOVA analysis of the dissolution data at 45 min for all tested brands showed that the dissolution behavior between different brands of glibenclamide tablet were significantly different at 95% CI (F > F crit; P< 0.05). Additionally, post ANOVA t-tests with Bonferroni correction were computed to point out which brands were not equivalent statistically with the comparator product with respect to their drug release profile at 95% CI. GL1, GL2, GL3 and GL6 were found to be significantly different in their drug release phenomena from comparator (GL5) product at 45 min (t stat > t crit).

The dissolution profiles of the generic and the innovator products at 8.5 pH phosphate buffer were presented in Fig 7. Since all tested brands exhibits drug release behavior of more than 75% at 30 minutes, it can be concluded that all tested glibenclamide tablets met USP-38 monograph requirements.

**Figure 7.**
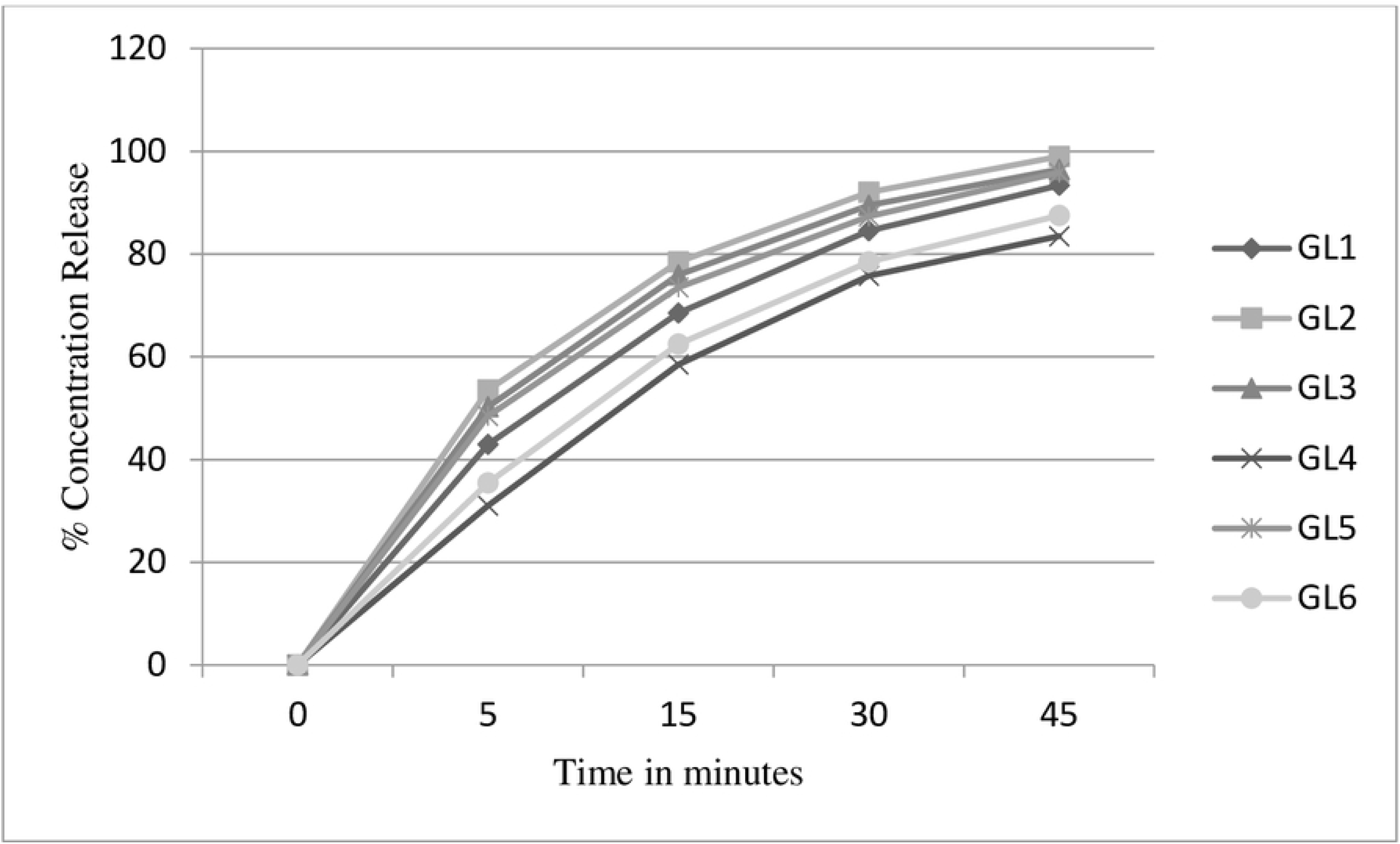
Dissolution profile of tested drugs at 8.5 pH phosphate buffer

The two way ANOVA analysis performed at 95% CI for the pharamacopoeially specified time, 30 minutes, found that there were significant differences in the release pattern of different glibenclamide brands (F > F crit; P < 0.05). As a result, post ANOVA t-tests with Bonferroni correction was done to observe the source of dissimilarity between comparator (GL5) and the tested generics at 95% CI. This comparative analysis showed that GL4 and GL6 were significantly different in their drug release pattern at 30 min when compared with the comparator product.

#### Dissolution profile evaluation

Model-independent approach of difference factor (*f*_*1*_) and similarity factor (*f*_*2*_) was computed for *in-vitro* dissolution profile studies to demonstrate the equivalence of all the generic glibenclamide tablets and the comparator product. [19] To ensure similarity and bioequivalence of two dissolution profiles, *f*_*1*_ should be between 0 and 15 whereas *f*_*2*_ should be between 50 and 100. [15-16, 19] In both cases, GL1, GL2 and GL3 were found to be similar in their dissolution behavior with comparator product (GL5) whereas GL6 was not. The similarity and dissimilarity values of tested products at pH of 7.5 and 8.5 were presented in Table 7. As a result, both *f*_*1*_ and *f*_*2*_ values justifies interchangeability of GL1, GL2 and GL3 generics with the comparator product. On the other hand, GL4 and GL6 might not be bioequivalent and therapeutically interchangeable with the comparator product (GL5).

**Table 7.** Similarity and dissimilarity factor comparison at pH of 7.5 and 8.5

| Product code | At pH of 7.5 | | At pH of 8.5 | | Status (Expected $f_2 > 50$ and $f_1 < 15$) |
| --- | --- | --- | --- | --- | --- |
| | $f_2$ | $f_1$ | $f_2$ | $f_1$ | |
| GL5 vs GL1 | 62.09 | 7.63 | 68.36 | 5.20 | Similar |
| GL5 vs GL2 | 79.95 | 2.99 | 66.80 | 5.82 | Similar |
| GL5 vs GL3 | 91.39 | 1.30 | 83.49 | 2.30 | Similar |
| GL5 vs GL4 | 42.32 | 16.41 | 45.24 | 13.78 | Dissimilar |
| GL5 vs GL6 | 48.25 | 14.44 | 48.90 | 13.51 | Dissimilar |

In order to ascertain the interchangeability of all products with the comparator product, drug release profiles of individual brands of Glibenclamide tablets were analyzed different kinetics models. The result of kinetic model studies for dissolution behaviors at pH 7.5 and 8.5 were presented in Table 8 and 9. The model that gives high correlation coefficient (r2) value is considered as the best fit model for the dissolution data. [16] Based on the findings, the Korsmeyer-Peppas model was found to best fit the dissolution data at both pH values. Therefore, it can be concluded that all brands under the investigation showed the similar drug release mechanism.

**Table 8.**
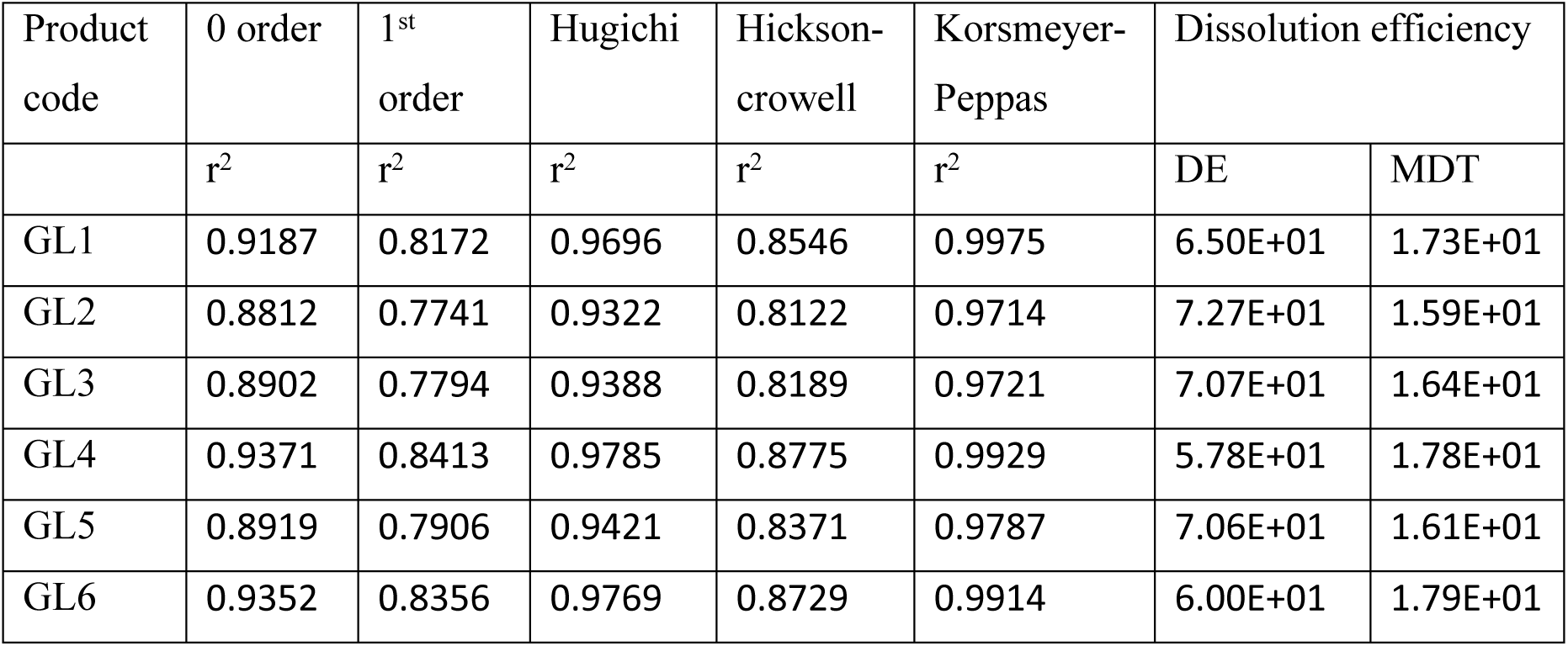
Correlation coefficient and DE values of kinetic models for dissolution data 7.5 pH

| Product code | 0 order | 1<sup>st</sup> order | Hugichi | Hickson-crowell | Korsmeyer-Peppas | Dissolution efficiency |  |
| --- | --- | --- | --- | --- | --- | --- | --- |
| | $r^2$ | $r^2$ | $r^2$ | $r^2$ | $r^2$ | DE | MDT |
| GL1 | 0.9187 | 0.8172 | 0.9696 | 0.8546 | 0.9975 | 6.50E+01 | 1.73E+01 |
| GL2 | 0.8812 | 0.7741 | 0.9322 | 0.8122 | 0.9714 | 7.27E+01 | 1.59E+01 |
| GL3 | 0.8902 | 0.7794 | 0.9388 | 0.8189 | 0.9721 | 7.07E+01 | 1.64E+01 |
| GL4 | 0.9371 | 0.8413 | 0.9785 | 0.8775 | 0.9929 | 5.78E+01 | 1.78E+01 |
| GL5 | 0.8919 | 0.7906 | 0.9421 | 0.8371 | 0.9787 | 7.06E+01 | 1.61E+01 |
| GL6 | 0.9352 | 0.8356 | 0.9769 | 0.8729 | 0.9914 | 6.00E+01 | 1.79E+01 |

**Table 9.**
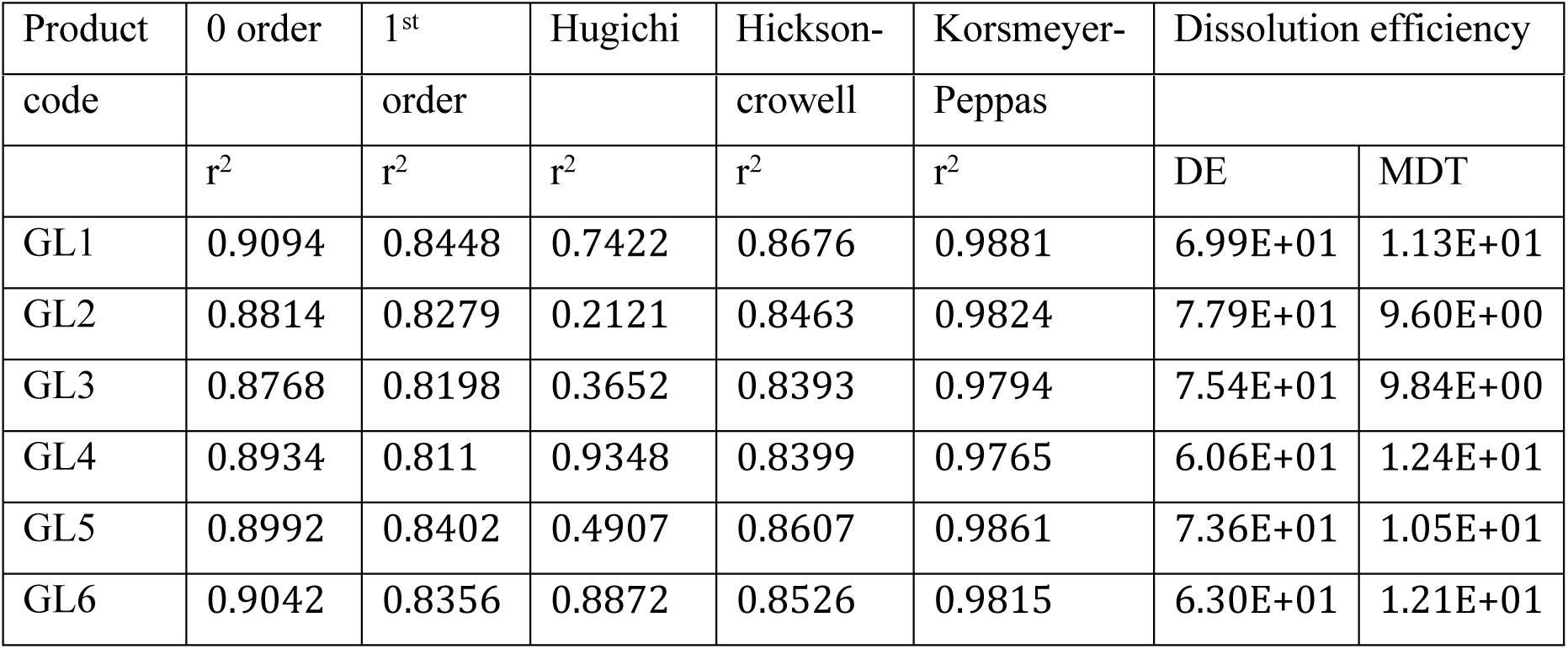
Correlation coefficient and DE values of kinetic models for dissolution data 8.5 pH

| Product | 0 order | 1<sup>st</sup> | Hugichi | Hickson- | Korsmeyer- | Dissolution efficiency |
| --- | --- | --- | --- | --- | --- | --- |

| code |  | order |  | crowell | Peppas |  |  |
| --- | --- | --- | --- | --- | --- | --- | --- |
|  | r<sup>2</sup> | r<sup>2</sup> | r<sup>2</sup> | r<sup>2</sup> | r<sup>2</sup> | DE | MDT |
| GL1 | 0.9094 | 0.8448 | 0.7422 | 0.8676 | 0.9881 | 6.99E+01 | 1.13E+01 |
| GL2 | 0.8814 | 0.8279 | 0.2121 | 0.8463 | 0.9824 | 7.79E+01 | 9.60E+00 |
| GL3 | 0.8768 | 0.8198 | 0.3652 | 0.8393 | 0.9794 | 7.54E+01 | 9.84E+00 |
| GL4 | 0.8934 | 0.811 | 0.9348 | 0.8399 | 0.9765 | 6.06E+01 | 1.24E+01 |
| GL5 | 0.8992 | 0.8402 | 0.4907 | 0.8607 | 0.9861 | 7.36E+01 | 1.05E+01 |
| GL6 | 0.9042 | 0.8356 | 0.8872 | 0.8526 | 0.9815 | 6.30E+01 | 1.21E+01 |

The Korsmeyer-Peppas equation (Log cumulative percentage of drug released versus log time) analysis confirm that both diffusion and erosion mechanism were involved in releasing the drug from the matrix tablets when values of drug release exponent (n) is in the range of 0.45 - 0.89. [15, 16] The calculated “n” values for all tested product falls within the range of 0.45 - 0.89 indicating that the release of the drug particle is due to both diffusion and erosion phenomena occurred throughout the course of dissolution study.

Moreover, the generics can be interchangeable for brand/innovator products when the difference between their dissolution efficiencies is within appropriate limits (±10%). [15, 20] Hence, it can be concluded that GL1, GL2 and GL3 had similar drug release profile with comparator (GL5) unlike GL4 and GL6 since their DE difference falls within the range of ±10%.

## Conclusions

This study evaluated the physicochemical quality and dissolution profile of six commercially available glibenclamide brands. The physical characteristic tests conducted on all tested product showed no deviation from the required specifications for hardness, friability, weight variation, package integrity and labeling completeness. All tested brands found to comply United States Pharmacopeia 38 monograph specification for their identification, label claim, content uniformity and dissolution. However, significant difference with respect to dissolution profile among tested brands GL4 and GL6 were confirmed with comparator product through model independent approach. Additionally, DE values were studied and confirmed that GL4 and GL6 were not therapeutically interchangeable with innovator product. These statistically significant differences in assay and dissolution profiles associated with the tested drugs are likely to reflect potential differences in clinical performance. Therefore, properly controlled bio-equivalence studies are strongly recommended to further investigate the potential problem that might associate with generic products.

## Acknowledgements

My profound and sincere gratitude go to Cadila Pharmaceutical Private Limited Company, Ethiopia for the generous donation of Working Standard and Reagents. I would like to extend my thanks to Jimma University Drug Quality Testing and Control Laboratory for providing necessary chemicals and reagents for this research.

